# Robust SARS-CoV-2-specific T-cell immunity is maintained at 6 months following primary infection

**DOI:** 10.1101/2020.11.01.362319

**Authors:** J Zuo, A Dowell, H Pearce, K Verma, HM Long, J Begum, F Aiano, Z Amin-Chowdhury, B Hallis, L Stapley, R Borrow, E Linley, S Ahmad, B Parker, A Horsley, G Amirthalingam, K Brown, ME Ramsay, S Ladhani, P Moss

## Abstract

The immune response to SARS-CoV-2 is critical in both controlling primary infection and preventing re-infection. However, there is concern that immune responses following natural infection may not be sustained and that this may predispose to recurrent infection. We analysed the magnitude and phenotype of the SARS-CoV-2 cellular immune response in 100 donors at six months following primary infection and related this to the profile of antibody level against spike, nucleoprotein and RBD over the previous six months. T-cell immune responses to SARS-CoV-2 were present by ELISPOT and/or ICS analysis in all donors and are characterised by predominant CD4+ T cell responses with strong IL-2 cytokine expression. Median T-cell responses were 50% higher in donors who had experienced an initial symptomatic infection indicating that the severity of primary infection establishes a ‘setpoint’ for cellular immunity that lasts for at least 6 months. The T-cell responses to both spike and nucleoprotein/membrane proteins were strongly correlated with the peak antibody level against each protein. The rate of decline in antibody level varied between individuals and higher levels of nucleoprotein-specific T cells were associated with preservation of NP-specific antibody level although no such correlation was observed in relation to spike-specific responses. In conclusion, our data are reassuring that functional SARS-CoV-2-specific T-cell responses are retained at six months following infection although the magnitude of this response is related to the clinical features of primary infection.

## Introduction

The SARS-CoV-2 pandemic has led to over 1 million deaths to date and there is an urgent need for an effective vaccine (1). There is considerable interest in understanding how adaptive immune responses act to control acute infection and provide protection from reinfection. Antibody responses against SARS-CoV-2 are characterised by responses against a range of viral proteins, including the spike, nucleocapsid and membrane proteins. A number of studies, however, have shown that the level of this antibody response declines over time and may even lead to loss of detectable virus-specific antibodies in a substantial proportion of individuals (2, 3). Information derived from study of immunity to related viruses such as SARS-CoV-1 and MERS (4) has shown that cellular immune responses against these viruses are maintained for much longer periods of time compared to antibody responses (5, 6). This has led to the hope that cellular responses to SARS-CoV-2 will similarly be of more prolonged duration (7, 8).

Studies to date have shown that virus-specific cellular responses develop in virtually all patients with confirmed SARS-CoV-2 infection (9). These responses remain detectable for several weeks following infection but it is currently unknown how they are maintained thereafter (10). In this study we characterised SARS-CoV-2-specific T cell immune responses in a cohort of 100 donors at 6-months post-infection.

## Results

### Characteristics of enrolled donors in the study

Blood samples were obtained from 100 convalescent donors at 6 months following initial SARS-CoV-2 infection in March-April 2020. Among the 100 donors, 77 (77%) were female and 23 (23%) were male with a median age of 41.5 years (22–65 years). None of the donors required hospitalisation at any time during the course of the study. 56 (45 female and 11 male) of the 100 donors who experienced clinical symptoms of respiratory illness were grouped as “symptomatic” and 44 (32 female and 12 male) who did not experience any respiratory illness were grouped as “asymptomatic”. There was no significant difference between the median age of the symptomatic (42.5 (23-61) years) and asymptomatic donors (40 (23-65) years).

### T-cell responses against SARS-CoV-2 are present in all donors and are 50% higher in donors with an initial symptomatic infection

Interferon gamma (IFN-g) ELISPOT analysis was used to determine the magnitude of the global SARS-CoV-2-specific T cell response. Peptide pools from a range of viral proteins, including spike, nucleoprotein and membrane protein, were used to stimulate fresh PBMC and the magnitude of the global SARS-CoV-2-specific T-cell response was determined. Median ELISPOT responses against the Spike glycoprotein (Spike); Nucleoprotein and Membrane (N/M); and ORF3a, ORF10, NSP8, NSP7A/b (Accessory) peptide pools were measured at 1 in 10,000 (0.010%), 12,500 (0.008%) and 66,666 (0.0015%) PBMC respectively (Figure 1A). Using the pre-2020 healthy donor PBMCs to set the cut-off point, 90 of 95 donors (95%) demonstrated a SARS-CoV-2-specific T-cell response to at least one protein with a median total value of 200 cells per million PBMC (1 in 5000) (Figure 1A). Eighteen donors did not have a demonstrable cellular response to Spike and no response to the N/M pool was seen in 9 individuals. No detectable response to any protein tested was seen in 5 donors by ELISPOT assay although all these donors responded by parallel intracellular cytokine analysis (Figure 1B).

**Figure 1.**
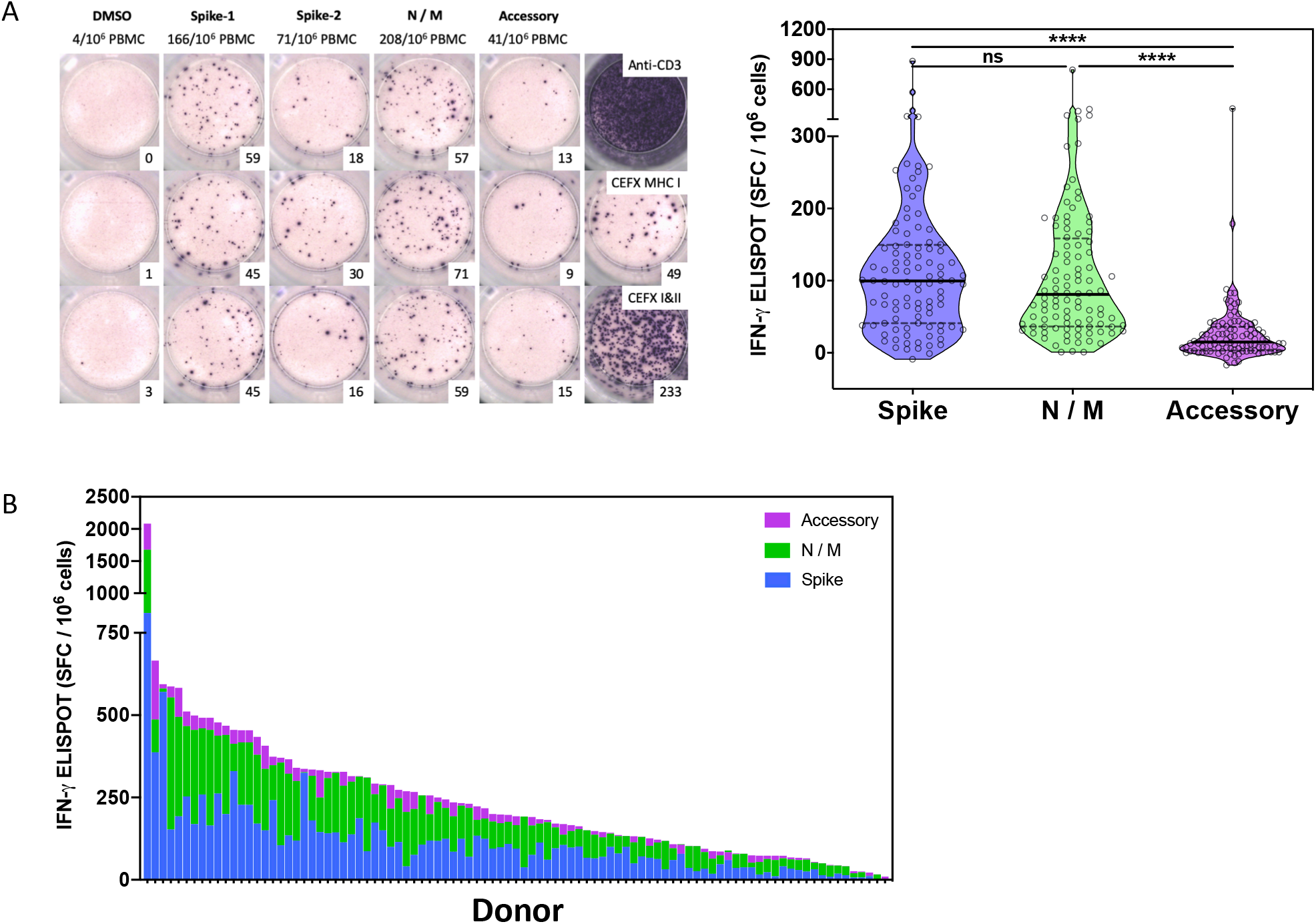
Robust T-cell immunity against SARS-CoV-2 is present in all donors at 6 months following primary infection. **A.** ELISPOT responses against SARS-CoV-2 protein pools at 6 months following primary infection. Left panel: A representative ELISPOT from 1 of 95 donors against Spike (pools 1 and 2), N/M and Accessory proteins (ORF3a, ORF10, NSP8, NSP7A/b), with DMSO as negative control and CEFX peptide pools and anti-CD3 as positive controls. Right panel: Summary data of all patients (N=95) studied according to Spike, N/M and Accessory peptide pools. Data in graph represented as SFC per million PBMC. Each point on violin plot represents a single donor. Bold black line represents median. The significance between pools was determined using a Friedman test with Dunn’s multiple comparison test, p < 0.0001(****). **B.** Aggregate ELISPOT response against SARS-CoV-2 proteins at 6 months following primary infection. The spot numbers were aggregated for individual donors and shown in a bar chart.

Considerable heterogeneity was observed in relation to the magnitude of this response. The global and peptide-specific responses were then assessed in relation to the clinical features at the time of primary infection. Importantly, median aggregate ELISPOT responses were 50% higher in donors who had initially demonstrated symptomatic disease compared to those with asymptomatic infection (Figure 2A). This profile was consistent against both spike and aggregate N/M proteins where values were 42% and 55% higher respectively in donors with initial symptomatic infection (Figure 2B). No association was seen between ELISPOT response and donor age.

**Figure 2.**
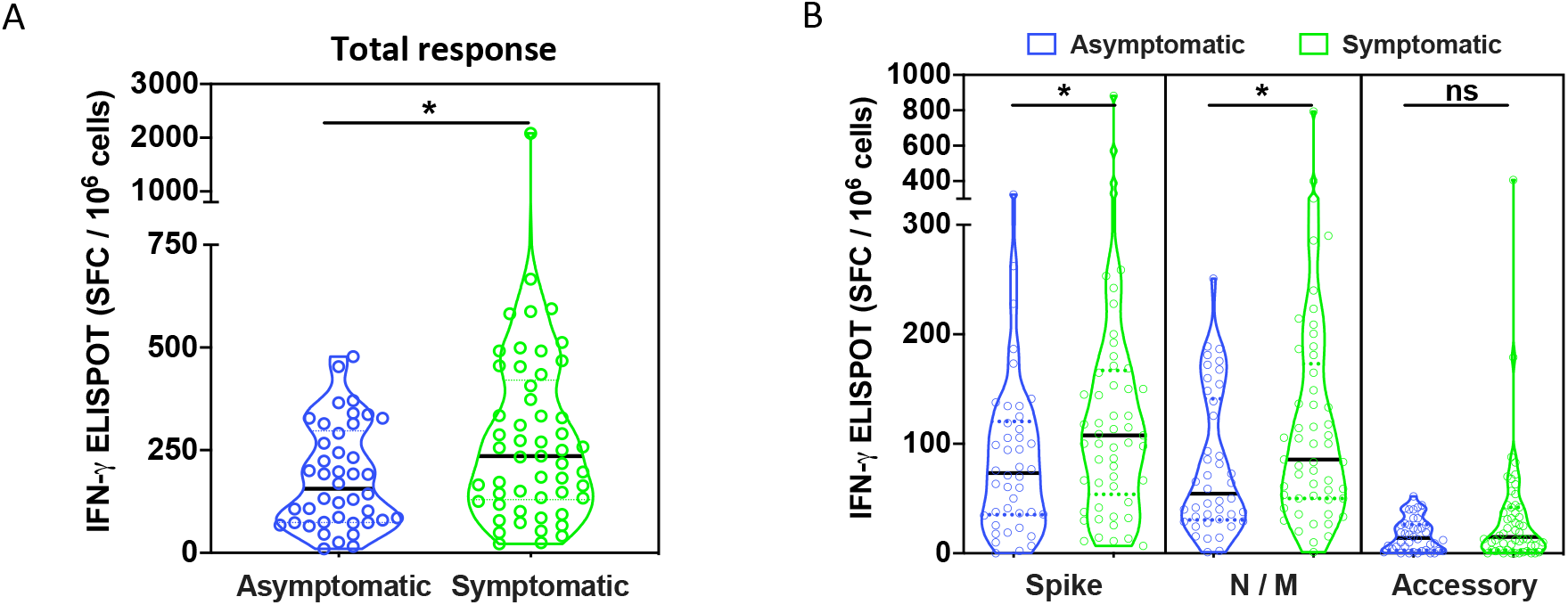
T-cell responses against SARS-CoV-2 are 50% higher in donors with an initial symptomatic infection. The cohort was divided into two groups according to symptoms at initial infection. **A**. The aggregated T-cell response (as SFC per million PBMC) against all peptide pools was compared between patients with (N=52) and without (N=43) respiratory symptoms. **B.** T-cell responses (as SFC per million PBMC) to Spike (pools 1 and 2), N/M, and Accessory proteins were compared between patients with and without symptoms. Each point on violin plot represents a single donor. Bold black line represents median. The significance was determined using Mann-Whitney testing, p < 0.05(*).

### SARS-CoV-2-specific T cell responses are characterised by a predominant profile of IL-2 production

Intracellular cytokine analysis was then utilised to assess the specificity and pattern of cytokine production from SARS-CoV-2-specific CD4+ and CD8+ T-cells in 100 donors. Virus-specific cytokine responses were seen in 96 people, including the 5 individuals that had been negative by ELISPOT analysis (Figure 3A). Interestingly, CD4+ virus-specific T cell responses were twice as frequent as CD8+ responses at this six-month time point (0.025% of CD4+ pool vs 0.012% of CD8+ pool respectively) (Figure 3B). In particular, mean CD4+ responses against spike and non-spike (N/M/Accessory) proteins were measured at 0.009% and 0.016% of the CD4+ repertoire respectively whilst corresponding values for CD8+ cells were 0.0045% and 0.0078% (Figure 3B). No differences were observed in the virus-specific CD4:CD8 ratio in relation to demographic factors such as age, symptomatic disease or gender.

**Figure 3.**
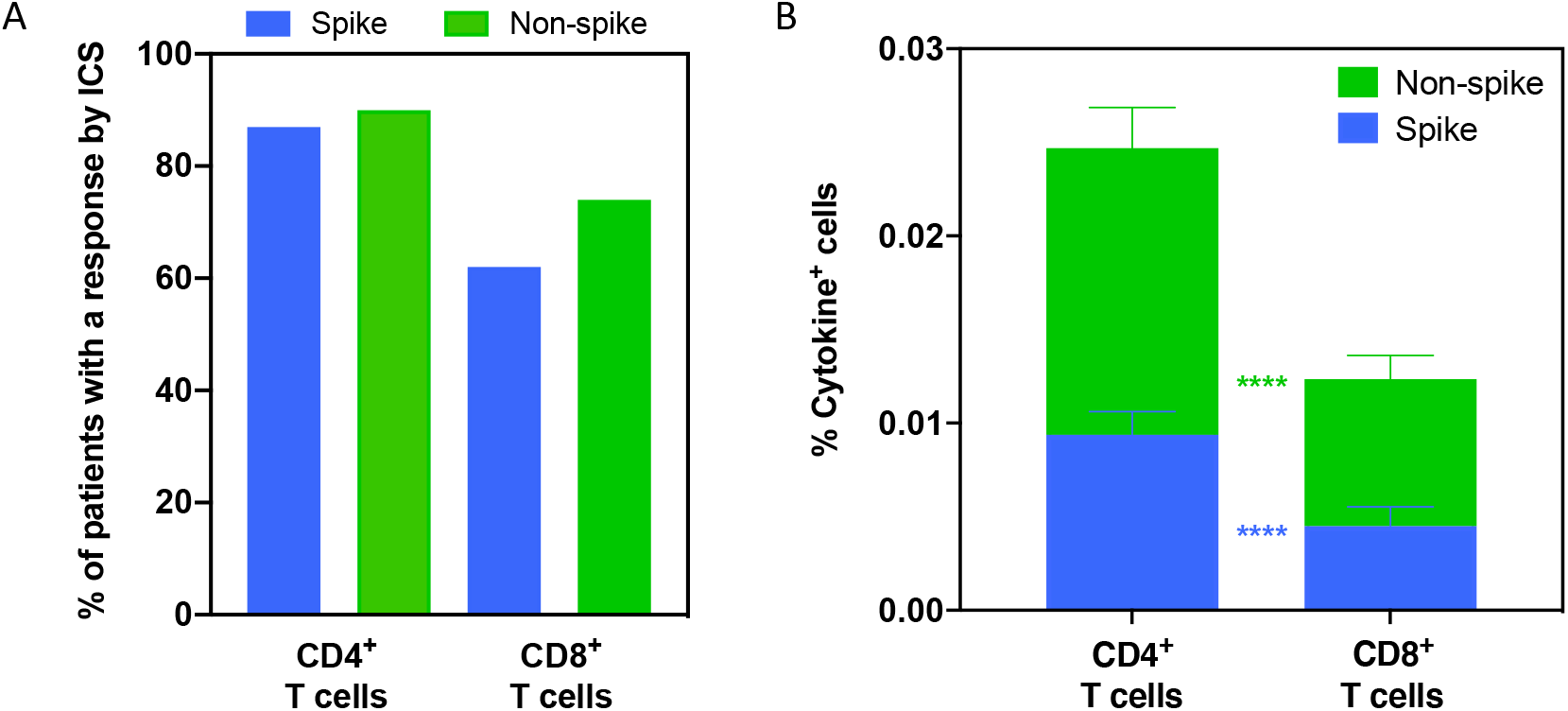
Overall detection of SARS-CoV-2-specific T cell responses by ICS. **A.** Proportion of donors (N=96) with a detectable IFN-y and/or IL-2 response by ICS for CD4+ T cells and CD8+ T cells against Spike and Nonspike proteins 6 months following primary infection. **B.** Aggregated IFN-y and IL-2 ICS responses for CD4^+^ and CD8^+^ T cells against Spike and Nonspike proteins. The significance was determined using Wilcoxon matched-pairs signed rank test, p < 0.001(****).

As expected, the pattern of cytokine production differed between the CD4+ and CD8+ subsets (Figure 4A, B). It was noteworthy that IL-2 responses were dominant within CD4+ subset (Figure 4B). Of note, the pattern of cytokine production by virus-specific CD4+ T cells was dependent on antigenic specificity. Single IFN-g, single IL-2 and dual positive IL-2+IFN-g+ T cells comprised 0.0016%, 0.0051% and 0.0027% of the spike-specific CD4+ T cell response respectively, compared to 0.0017%, 0.0105% and 0.0031% of the non-spike-specific repertoire (Figure 4C).

**Figure 4.**
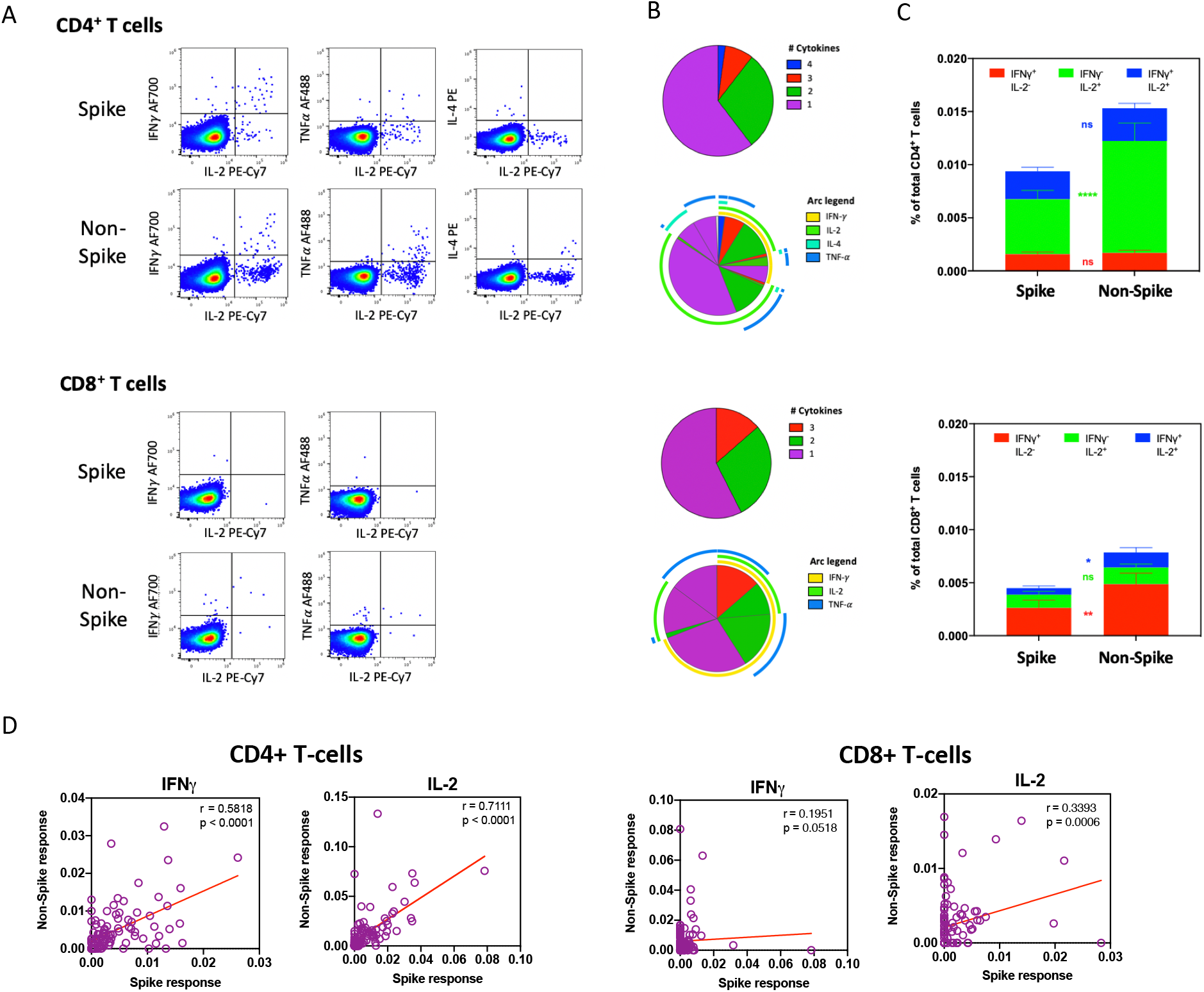
SARS-CoV-2-specific T-cell responses are characterised by a predominant profile of IL-2 production. **A.** Features of CD4+ and CD8+ T-cell responses against SARS-CoV-2 proteins by intracellular cytokine staining at 6 months. Representative flow plots of CD4+ (Top panel) and CD8+ (Bottom panel) T-cell responses against peptide pools from Spike or non-spike (aggregate of N, M, ORF3a, ORF10, NSP8 and NSP7A/b) proteins. **B.** Polyfunctional analysis of SARS-CoV-2-specific CD4+ and CD8+ T-cells at 6 months. Relative distribution of single or multiple cytokine responses in CD4+ (Top panel) and CD8+ (Bottom panel) T-cells, and pattern of co-expression of IL-2, IFN-γ, TNF-a and IL-4 in SARS-CoV-2-specific T-cells. **C.** Aggregate ICS responses for CD4+ and CD8+ T-cells against Spike and Non-spike proteins according to IFN-γ and/or IL-2 production. The significance was determined using Wilcoxon matched-pairs signed rank test, p < 0.05(*), p < 0.01(**), p < 0.0001(****). **D.** Correlation of Spike and Non-spike responses according to IFN-γ and IL-2 production by CD4+ (left panel) and CD8+ (right panel) T-cells at 6 months. Spearman’s Rank correlation was used to test the significance, and p value and r value (correlation coefficient) are indicated for each panel.

Analysis of the Th cytokine profile in the supernatants of overnight *ex vivo* peptide-stimulated ELISPOT cultures confirmed IL-2 to be the dominant cytokine released by SARS-CoV-2 specific T cells with variable TNFα release, alongside IFN-g as detected by ELISPOT. There was no release of cytokines indicative of other Th subsets including Th2 and Th17 (Supplementary Figure).

The magnitude of CD4+ T cell responses against spike and non-spike proteins within each individual was strongly correlated (Figure 4D). However, this association was less marked for the CD8+ subset where responses were dominant against non-spike proteins (Figure 4D).

### The magnitude of the T-cell response at six months correlates with peak antibody level and reduced rate of antibody waning against nucleoprotein

We next assessed how the magnitude, phenotype and cytokine profile of the virus-specific cellular immune response at six months correlated with the prospective profile of antibody production in the six months since infection. Antibody levels against both the Spike glycoprotein and nucleoprotein were available at serial time points from all donors (Figure 5A). These were used to define both the peak value of antibody level against each protein and the rate of decline in antibody level over the subsequent two months. Peak antibody levels against both spike and nucleoprotein were observed typically at the second month of sampling (Figure 5A). Antibody levels fell by approximately 50% during the two months after peak level but stabilised somewhat thereafter although spike-specific responses continued to decline (Figure 5A).

**Figure 5.**
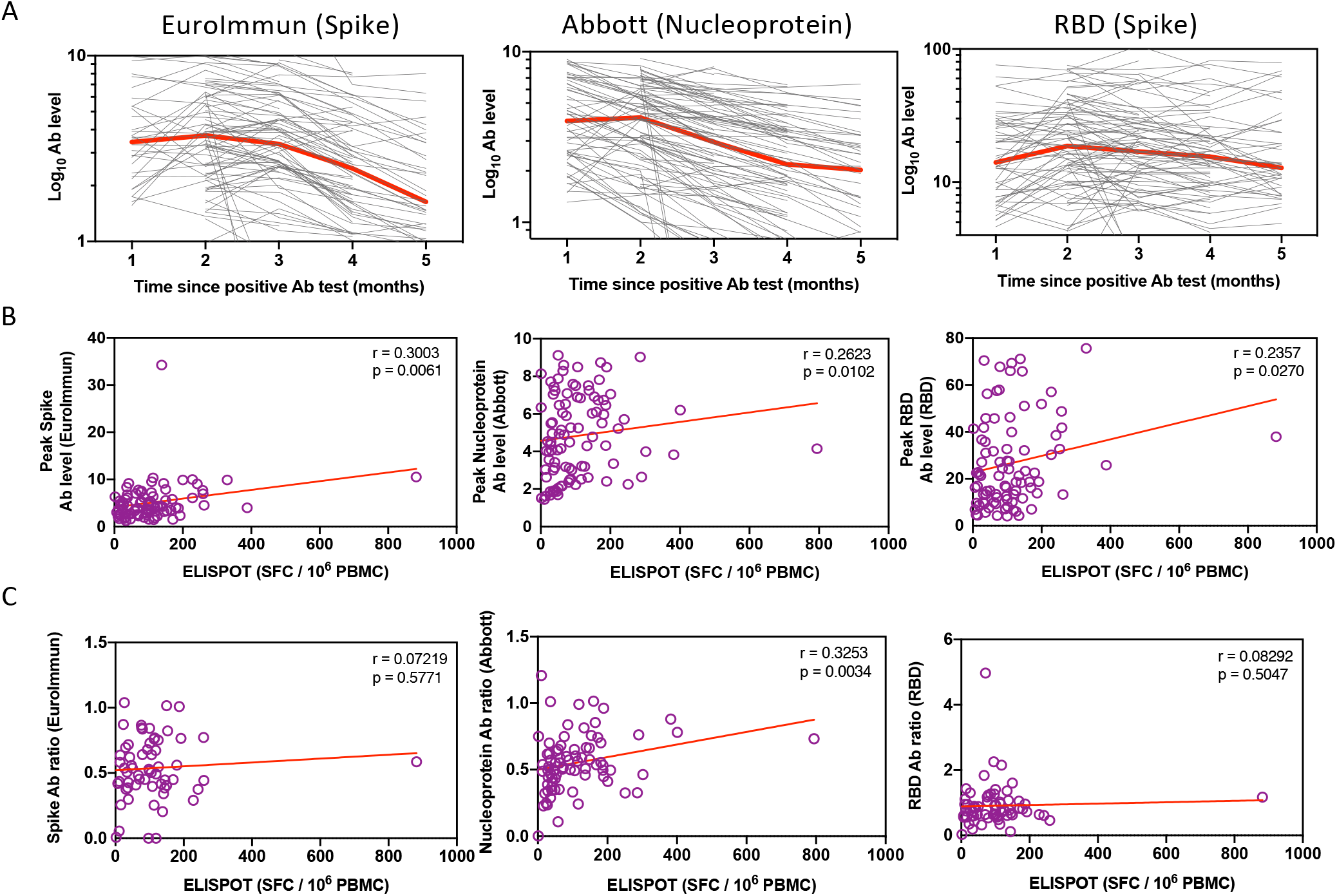
The magnitude of the T-cell response at six months correlates with peak antibody level and reduced rate of antibody waning against nucleoprotein. **A**. Antibody levels against Spike(N=81), Nucleoprotein(N=94), and RBD(N=87) of all patients at each time point post infection were plotted. Each grey line represents an individual patient. The median antibody level over time is shown in red. **B.** The correlation of ELISPOT responses at 6 months against peak antibody levels (Spike: N=82, Nucleoprotein: N=94, and RBD: N=87) were assessed for each antibody. **C.** Correlation of ELISPOT responses at 6 months with rate of antibody decline (expressed as ratio of ‘antibody level at 2 months after peak level’: ‘antibody peak level’) (Spike: N=60, Nucleoprotein: N=79, and RBD: N=67). The line represents linear regression. Spearman’s Rank correlation was used to test the significance, and p value and r value (correlation coefficient) are indicated in each panel.

Interestingly, the magnitude of the T cell ELISPOT response at 6 months against the spike protein was strongly correlated with magnitude of the peak antibody level against both spike protein and the RBD domain (Figure 5B). A similar correlation was observed between the cellular response to the N/M pool and the peak level of N-specific antibody (Figure 5B).

The rate of antibody decline was then assessed in relation to the profile of the cellular immune response at 6 months. Relative preservation of the N-specific antibody response was seen in donors with stronger N and M-specific T cell responses at six months suggesting the cellular responses may act to support antibody production (Figure 5C). However, no such association was observed in relation to spike-specific responses.

Finally, we also assessed the profile of CXCR5 expression on virus-specific T cells and related this to the pattern of stability of the virus-specific antibody response as positive correlations have been observed previously in HIV infection (11). High numbers of circulating Tfh CD4+ T cells have been seen in severe acute infection (12) but at 6 months CXCR5 was expressed on only 7% of virus-specific CD4+ T cells and no correlation was observed with the profile of Ab level following infection.

## Discussion

The magnitude and quality of the immune memory response to SARS-CoV-2 will be critical in preventing reinfection. Here we undertook, to our knowledge, the first assessment of the SARS-CoV-2-specific T cell immune response at six months following primary infection in a unique cohort of healthy adults with asymptomatic or mild-to-moderate COVID-19.

The major finding was that virus-specific T cells were detectable in all donors at this extended follow-up period. A high prevalence of detectable T-cell immunity has been described in studies performed at earlier time points after infection and our findings indicate that robust memory is maintained for at least 6 months. Approximately 1 in 4000 PBMC were SARS-CoV-2-specific which is broadly comparable to findings within the first three months after infection. These values are lower than typical responses against persistent herpesviruses (13) but comparable to those against acute respiratory viruses, including SARS-CoV (14, 15).

The magnitude of the T cell response was heterogeneous and may reflect the reports of remarkable diversity in the profile of the T cell immune response during acute infection (16). A striking feature was that the magnitude of cellular immunity was considerably higher in donors who had experienced symptomatic disease. Indeed, median ELISPOT responses were 50% higher within this group and demonstrate that the initial relative ‘setpoint’ of cellular immunity established following acute infection is maintained for at least 6 months. A similar pattern has been observed within the first few weeks following acute SARS-CoV-2 infection in patients recovering from severe versus mild disease (17) and also in patients after SARS infection (30). This is likely to reflect a response to higher viral loads and inflammatory mediators during acute infection (18, 19) although it is also possible that an elevated adaptive immune response during primary infection can itself act as a determinant of the clinical phenotype (20). Cellular responses have a direct protective effect against severe coronavirus infection (21) and also support antibody production. Indeed, cytokine analysis showed that the CD4+IL-2+ subset was most significantly elevated in the symptomatic group. The finding of lower levels of T cell immunity in asymptomatic donors at 6 months after infection might potentially add to concerns that this group may be more susceptible to later re-infection. However, it is also possible that the quality of the T cell response at the time of initial infection was sufficient to provide clinical protection. The relative susceptibility of patients with initial asymptomatic disease to episodes of re-infection, either clinically silent or symptomatic, will therefore need to be assessed over time.

It was noteworthy that CD4 T cells responses against SARS-CoV-2 outnumbered CD8 effector cells by ratio of 2 to 1. Again, a similar pattern has been demonstrated at earlier time points after SARS-CoV-2 infection and may reflect high levels of viral protein uptake by antigenpresenting cells and cross presentation to the CD4+ positive T-cell pool or preferential expansion of CD4+ T cells (22). Furthermore, cytokine analysis showed that IL-2 was the major cytokine produced by virus-specific CD4+ cells, indicating a proliferative potential which may auger well for long-term immune memory (23). IFN-g responses are broadly equivalent to IL-2 at early time points after infection (24) but the profile at 6 months suggests that the relative proportion of Th1 effector cells decreases over time or they revert to central memory state (25). Polyfunctional T cells are typically associated with superior pathogen control (26) and studies on SARS-CoV-2 infections have revealed decreased cytokine functionality in patients with severe disease (17). The majority of CD4+ T cells at six months expressed only a single cytokine and production of three or four cytokines was observed in <15% of cells. These results were consistent with comprehensive analysis of the cytokine profile released by SARS-CoV-2 specific T cells in supernatants from the ELISPOT assay which showed that IL-2 was consistently the dominant cytokine. Interestingly, low levels of IL-10 were released in response to all the peptide pools and as IL-10 production has been reported by subsets of murine Influenza and Coronavirus specific T cells (27, 28) these represent an interesting population of cells for future investigation. Low and variable concentrations of IL-4 and TNFα were also detected.

Of note, the pattern of cytokine production by CD4+ T cells varies with protein specificity, as seen in earlier reports (17). Single IL-2 or IFN-g producing cells were predominant against both spike and structural proteins but the former population was significantly greater in the CD4+ response against non-spike proteins, indicating that a retained Th1 effector profile is more common within spike-specific T cell pool. These dual-positive populations are associated with elite immunological control of HIV infection and indicate that spike-specific T cells preferentially retain characteristics of both effector function and proliferative potential *in vivo*.

The expression of CXCR5 on CD4+ T cells has been correlated with the magnitude and persistence of humoral immunity in the setting of HIV infection (29). We, however, observed that CXCR5 was expressed on only 7% of virus-specific CD4+ T cells suggesting that circulating virus-specific follicular helper cells are not sustained after infection and no clear relationship was noted between CXCR5 expression and either the magnitude or the rate of decline of SARS-CoV-2-specific antibody level. Findings in acute infection have also failed to correlate cTFh frequencies with the plasmablast response and suggest that non-CXCR5+ CD4+ T cell help may also operate (16).

One of the valuable features of our cohort was the availability of monthly antibody levels against the spike and nucleoproteins in the first six months after infection. The finding that that higher T cell responses at 6 months against N/M proteins correlated with slower decline in N-specific antibody levels indicates that vaccine approaches that elicit strong cellular immune responses against this protein are likely to be valuable for sustaining stable antibody responses. In contrast, T-cell responses against Spike were not related to the rate of decline of antibodies against that protein. Nevertheless, spike protein-specific cellular responses were present in >80% of individuals at 6 months after mild to moderate infection and are also recognised as an immunodominant protein following SAR-CoV-1 infection (30). Spike glycoprotein is the major immunogen used in current vaccine trials and these findings indicate that strong and sustained spike-specific T-cell immunity is likely to be required to sustain immune protection and should be assessed in analysis of optimal vaccine strategies. Our finding that T cell responses against M/N proteins are equally as high as Spike responses at 6-months after natural infection suggest that these proteins could also represent valuable components of future vaccine strategies.

Our findings demonstrate that robust cellular immunity against SARS-CoV-2 is likely to be present within the great majority of adults at six months following asymptomatic and mild-to-moderate infection. These features are encouraging in relation to the longevity of cellular immunity against this novel virus and are likely to contribute to the relatively low rates of reinfection that have been observed to date (31). Further studies will be required to assess how these immune responses are maintained over the longer term.

## Acknowledgements

This work was partly funded by UKRI/NIHR through the UK Coronavirus Immunology Consortium (UK-CIC). KV is supported by Blood Cancer UK (grant 17009) and AD is supported by MRC (grant MR/R011230/1).

This research was carried out with the support of the NIHR Manchester Clinical Research Facility. BP and AH are supported by the NIHR Manchester Biomedical Research Centre.

The views expressed are those of the author(s) and not necessarily those of the NHS, the NIHR or the Department of Health.

## Materials and Methods

### Ethical Statement and Clinical definitions

This study was approved by PHE Research Support and Governance Office (R&D REGG Ref NR 0190). Donors were recruited from a cohort of staff at Public Health England (PHE) that has been monitored for acute infection in March-April 2020. Written informed consent was obtained from all donors. The majority of donors were asymptomatic at the time of initial infection and none were admitted to hospital. All donors were SARS-CoV-2 seropositive using either the Euroimmun, RBD or Abbott test. Serum samples were taken at monthly intervals and assessed by the Euroimmun anti-spike ELISA or the Abbott anti-N assay system. Mean log values were used to determine antibody levels. Cut off levels for positivity were set at >0.8 for the Abbott (N), >5 for RBD (S) and >1.1 for EuroImmun (S) assay as described earlier(32). Blood samples for cellular analysis were taken at 6 months from the initial PCR-positive test and SARS-CoV-2 sero-negative and pre-2020 healthy donor samples were used as controls.

### Synthetic peptides

Pepmixes of 15-mer peptides overlapping by 11 amino acid residues covering the major proteins of SARS-CoV-2 (Spike Glycoprotein (PM-WCPV-S), Membrane protein(PM-WCPV-VME), nucleoprotein (PM-WCPV-NCAP), ORF3A (PM-WCPV-ORF3A),ORF 7A/B(PM-WCPV-NS7A/7B), ORF10 (PM-WCPV-ORF10) and Non-Structural protein 8 (PM-WCPV-NS8) (JPT Peptide Technologies, Berlin, Germany). A pool of immuno-dominate epitopes from common viruses including Cytomegalovirus (CMV), Epstein-Barr virus (EBV) and influenza virus (PM-CEFX) was included as positive control (JPT Peptide Technologies).

### ELISPOT assay

T cell responses were assessed by ELISPOT assay using a Human IFNγ ELISPOTPro kit (Mabtech, NS, Sweden) following the manufacturer’s instructions. Briefly, freshly isolated PBMC were rested overnight prior to assay. Plates were washed with filtered PBS (Sigma Aldrich, Missouri, US), and blocked with culture media containing 10% batch tested FBS (Gibco, Thermo Fisher Scientific, Massachusetts, US). As standard 3×10^5^ PBMC per well were stimulated in triplicate with overlapping peptide pools (JPT Peptide Technologies) at a concentration of 1ug/ml of individual peptide for 18hrs. In some cases, assays were run in duplicate or with 2.5×10^6^ PBMC as a minimum. Negative controls comprising DMSO and positive controls, anti-CD3 and CEFX pepmix (JPT Peptide Technologies), were also included. Spots were counted using an AID ELISPOT Reader System (AID GmbH, Strasberg, Germany). Mean spot counts for negative control wells were subtracted from the mean of test wells to generate normalised readings, these are presented as Spot Forming Cells per million input PBMC (SFC/10^6^ PBMC).

### Intracellular cytokine staining (ICS)

Freshly isolated PBMC were rested overnight prior to the assay. 1.5×10^6^ PBMCS were stimulated with peptide pools for Spike, or a combination of NCAP, VME1, ORF10, NS7A, NS7B, AP3A, and NS8 at a concentration of 1ug/ml of individual peptide for 6hrs in the presence of protein transport inhibitor cocktail (Ebioscience, San Diego, CA, US). After incubation, PBMCs were harvested and washed before adding fixable red viability dye (Thermo Fisher Scientific) and cell surface antibodies anti-CD3-PerCP5.5 (Biolegend, San Diego, CA, US), anti-CD4-APC-Cy7(Biolegend), anti-CD8-BV510 (Biolegend), anti-PD-1-Pacific Blue (Biolegend) and anti-CXCR5-APC (Biolegend). Staining was performed at 4°C for 30 mins. Then the PBMCs were washed and fixed with Ebioscience IC Fixation buffer at 4°C overnight. Following incubation, the fixed cells were permeabilized with 0.1% Triton–X 100 at (Sigma Aldrich) on ice for 30min, washed with PBS and stained at 4°C for 50 mins with the intracellular antibodies, anti-TNF-α-FITC (Biolegend), anti-IFN-g-AF700 (Biolegend), anti-IL-2-PE-Cy7 (Biolegend) and anti-IL-4-PE (Biolegend). All antibodies were purchased from Biolegend. Finally, the cells were washed in PBS before analysis on Beckman Coulter Gallios Flow cytometer (Beckman Coulter, High Wycombe, US). Negative controls without peptide-stimulation were also included for each donor sample. Flow cytometry data was analysed using FlowJoTM v.10 software (FlowJo LLC, Ashland, Oregon, US). Pestle and SPICE software (version 6) was used to determine the frequency of different cytokine response patterns based on all possible combinations (33).

### Supernatant Cytokine Profile

Following overnight peptide stimulation in ELISPOT assays 50ul of supernatant was removed and combined from two duplicate wells and cryopreserved at −80°C. Supernatant from eleven donors responding in the ELISPOT assay were profiled using a 12-plex Legendplex T Helper Cytokine Panel Version 2 (Biolegend) following the manufactures instructions. Cytokine beads were analysed on a BD LSR II flow cytometer (BD Biosciences, San Jose, CA, US). Data was analysed with Legendplex Software (Biolegend) and the average cytokine level determined from two duplicate samples.

### Statistical analysis

Statistical analysis was performed with GraphPad Prism 8. A Mann-Whitney 2-tailed U test was used to compare variables between two groups, a Wilcoxon matched-pairs signed rank test was used to compare paired non-parametric data, and a Friedman test with Dunn’s multiple comparisons test was used to compare non-parametric data between more than 2 groups. Correlations were performed via Spearman’s rank correlation coefficient. Two-way ANOVA with Dunnett multiple comparisons test was used to determine significance of cytokine profile data. Statistical significance was determined as *P<0.05, **P<0.01, ***P<0.001 and ****P<0.0001.

## Notes

### Competing Interest Statement

The authors have declared no competing interest.

