## supplementary figure for "Robust SARS-CoV-2-specific T-cell immunity is maintained at 6 months following primary infection"

**Supplementary figure: Characterisation of the Th cytokine profile released by SARS-CoV-2 specific cells after peptide stimulation**.

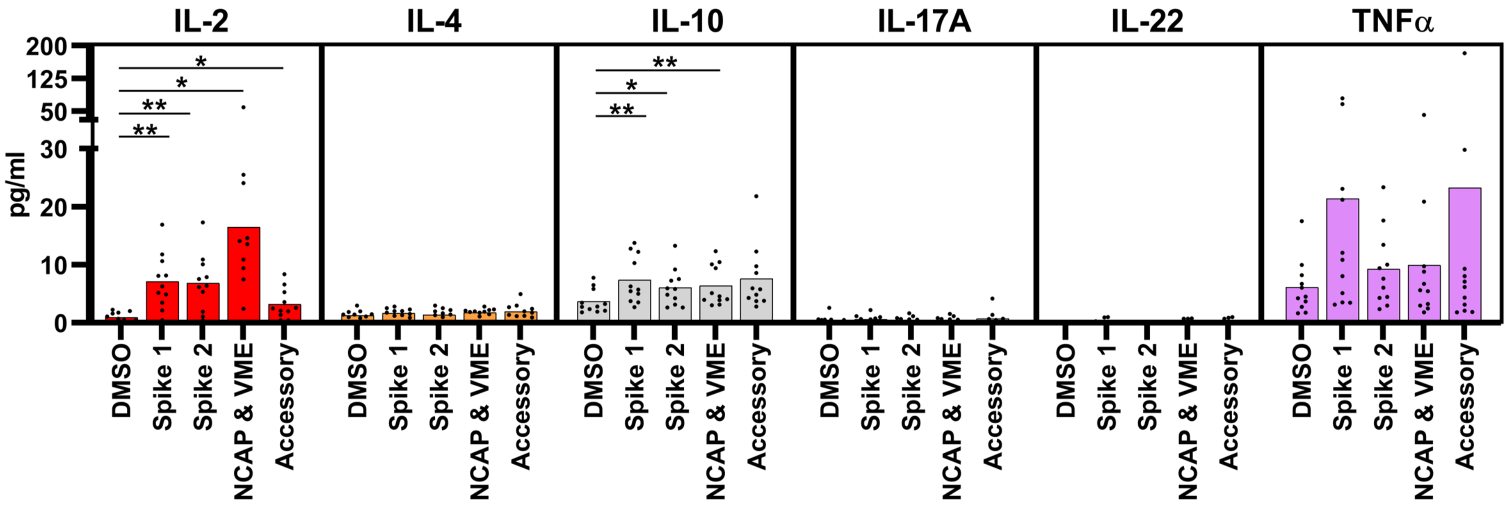


**Supplementary figure:** Characterisation of the Th cytokine profile released by SARS-CoV-2 specific cells during peptide stimulation shows IL-2 is consistently the dominant cytokine released. Supernatant from the wells of ELISPOT assays from eleven responding donors was analysed to assess the release of cytokines representative of classical Th subsets. In addition to the shown, IL-5, -9, -13 and IL-17F were not detected. Two-way ANOVA with Dunnett multiple comparisons test, * $p < 0.05$, ** $p < 0.01$.
